# Reward Memories Bias Instrumental Rule Selection via the Orbitofrontal-to-Secondary Motor cortex Pathway

**DOI:** 10.64898/2026.08.29.748039

**Authors:** Margo Le, Jessica Milmoe, Lydia Nungesser, Amanda Maheras, Ronald Keiflin

## Abstract

Reward-predictive cues can influence decision-making and promote the pursuit of specific outcomes. This influence is classically studied using specific Pavlovian-to-instrumental transfer (sPIT), in which Pavlovian cues bias action selection and promote the instrumental action (often a left or right lever press) directed toward the cued outcome. However, naturalistic reward pursuit often extends beyond selecting discrete actions and requires selecting abstract rules that organize multiple actions into goal-directed sequences. The influence of cued reward memories on such rule selection has received less attention, and the neural circuits mediating this effect remains largely unknown. Here, we developed the specific Pavlovian-to-Rules-to-Instrumental Transfer (sPRInT) task, an adaptation of sPIT designed to examine how Pavlovian reward cues bias instrumental rule selection. We then used pathway-specific chemogenetic silencing to test the contribution of orbitofrontal cortex projections to secondary motor cortex (OFC→M2) to this effect. Rats expressing hM4Di or mCherry in OFC→M2 neurons learned a three-step instrumental sequence (sample lever – nosepoke – choice lever). In alternating blocks, rats used either a delayed non-match-to-sample rule or a visually guided rule to select the final action in the sequence and earn distinct outcomes (dNMTS–O1; VIS–O2). In a second phase, two distinct auditory cues were established as Pavlovian predictors of the two outcomes (S1–O1; S2–O2). Finally, in nonrewarded probe tests preceded by DCZ injections, we examined how presentation of these Pavlovian cues biased instrumental choices by promoting the use of an abstract rule. In control (mCherry) rats, Pavlovian cues promoted the adoption of the rule corresponding to the cued outcome (S1 promoted dNMTS; S2 promoted the VIS rule). In hM4Di rats, DCZ-mediated inactivation of OFC→M2 projection neurons diminished this effect. In contrast, OFC→M2 inhibition spared conventional sPIT, in which reward-predictive cues directly biased action selection without requiring abstract rules or extended action sequences. These findings demonstrate that cued reward memories can govern abstract rule selection and identify OFC→M2 as a critical circuit for translating those memories into goal-appropriate behavioral strategies.

## INTRODUCTION

Imagine this: as you walk home, debating between pizza and sushi for dinner, you smell the enticing aroma of wood-fired pizza wafting from your local pizzeria. That settles it —pizza it is! This simple scenario highlights the powerful influence of reward-predictive cues on decision-making. Specifically, by evoking the memory of certain outcomes, reward-predictive Pavlovian cues can bias decision-making and promote the instrumental actions directed at those outcomes (Walker, 1942; Estes, 1948; Rescorla and Solomon, 1967; Kruse et al., 1983; Balleine and Ostlund, 2007; de Wit and Dickinson, 2009; Allman et al., 2010; Holmes et al., 2010; Cartoni et al., 2016; Mahlberg et al., 2021; Badioli et al., 2024). This interaction between Pavlovian and instrumental processes is highly adaptive and ensures that reward-seeking actions are efficiently distributed and aligned with available outcomes. Dysregulation of this process can lead to impaired decision-making and maladaptive motivated behaviors and is observed across several neuropsychiatric disorders (Hogarth et al., 2013; Watson et al., 2014; Garbusow et al., 2022; Chen et al., 2023).

Experimentally, the influence of stimulus-evoked predictions on action selection can be precisely assessed using the specific Pavlovian-to-Instrumental Transfer (sPIT) paradigm. This paradigm typically consists of three phases. In the Pavlovian phase, two different stimuli are established as predictors of two different outcomes (S1–O1; S2–O2). In the instrumental phase, subjects learn that two different actions—often a left or right lever press—produce these respective outcomes (A1–O1; A2–O2). Finally, the transfer test measures the influence of Pavlovian stimuli on instrumental behavior by presenting the stimuli while subjects are given the opportunity to perform either of the two instrumental actions. A specific PIT effect is observed when Pavlovian stimuli selectively enhance the instrumental action associated with the predicted outcome (i.e., S1 promotes A1, and S2 promotes A2).

By separating the Pavlovian and instrumental learning phases, the sPIT paradigm prevents the development of direct stimulus-response associations, ensuring that reward-predictive cues influence instrumental behavior via the activation of outcome-specific memories. For this reason, sPIT has become the gold standard for assessing the impact of cues on reward-guided decision making and has facilitated key discoveries in the neural circuits of motivated behaviors (Corbit and Balleine, 2005; Corbit and Janak, 2007; Bray et al., 2008; Talmi et al., 2008; Laurent et al., 2014; Leung and Balleine, 2015; Lichtenberg et al., 2017; Sias et al., 2021; Malvaez et al., 2026)

However, one limitation of sPIT is that it fails to account for the fact that, beyond specific discrete actions (e.g., left or right lever press), naturalistic reward pursuit typically involves multiple actions organized into logical sequences (Lashley, 1951; Cooper and Shallice, 2000; Botvinick, 2008; Grafton et al., 2009). The organization of discrete actions within these sequences is often guided by goal-specific, higher-order rules, which dictate the environmental features and cognitive strategies that are relevant for the pursuit of the expected outcome. While sPIT assesses the influence of reward cues on *action selection*, the influence of reward cues on *rule selection* has received less attention, and the neural circuits underlying this process remain largely unknown.

Candidate regions for this cue-evoked rule selection process include the orbitofrontal cortex (OFC) and the secondary motor cortex (M2). The OFC plays a critical role in encoding predictive relationships between events and generating outcome-specific predictions (i.e. sensory-rich representations of an anticipated outcome’s identity) (Rolls, 2004; McDannald et al., 2011; Stalnaker et al., 2014; Wilson et al., 2014; Howard et al., 2015; Costa et al., 2023). M2 contributes to the representation of instrumental rules and the planning and coordination of complex action sequences (Gremel and Costa, 2013; Barthas and Kwan, 2017; Ohbayashi, 2021; Sánchez-Fuentes et al., 2021; Schreiner et al., 2022; Augusto et al., 2025). Critically, the OFC sends substantial projections to M2, and this OFC→M2 pathway undergoes structural plasticity during rule learning (Hoover and Vertes, 2011; Bedwell et al., 2014; Johnson et al., 2016). Collectively, these studies suggest that the influence of cue-evoked outcome memories on rule selection might be mediated by OFC projections to M2.

To test this hypothesis we designed a novel rodent behavioral paradigm: the specific Pavlovian-to-Rules-to-Instrumental Transfer (sPRInT) paradigm. Adapted from sPIT, this paradigm replaces discrete action-outcome contingencies (A1–O1; A2–O2) with sequence rule contingencies (Rule1–O1; Rule2– O2). Using the sPRInT paradigm in combination with chemogenetic inactivation of the OFC→M2 pathway, we demonstrate that cue-evoked outcome memories bias instrumental rule selection, and that this effect relies on the functional integrity of the OFC→M2 pathway.

## RESULTS

### Cued reward memories bias abstract rule selection, and this effect is reduced by OFC→M2 inactivation

To gain control over OFC→M2 pathway and examine its contribution to abstract rule selection in reward pursuit, we used an intersectional viral strategy to express the inhibitory DREADD hM4Di, or the control transgene mCherry, selectively in OFC neurons projecting to M2 in male and female rats (**Figure 1A-B**). DCZ-mediated chemogenetic inhibition of the OFC-M2 pathway was then performed only during the transfer tests following instrumental and Pavlovian learning (**Figure 1C)**.

**Figure 1.**
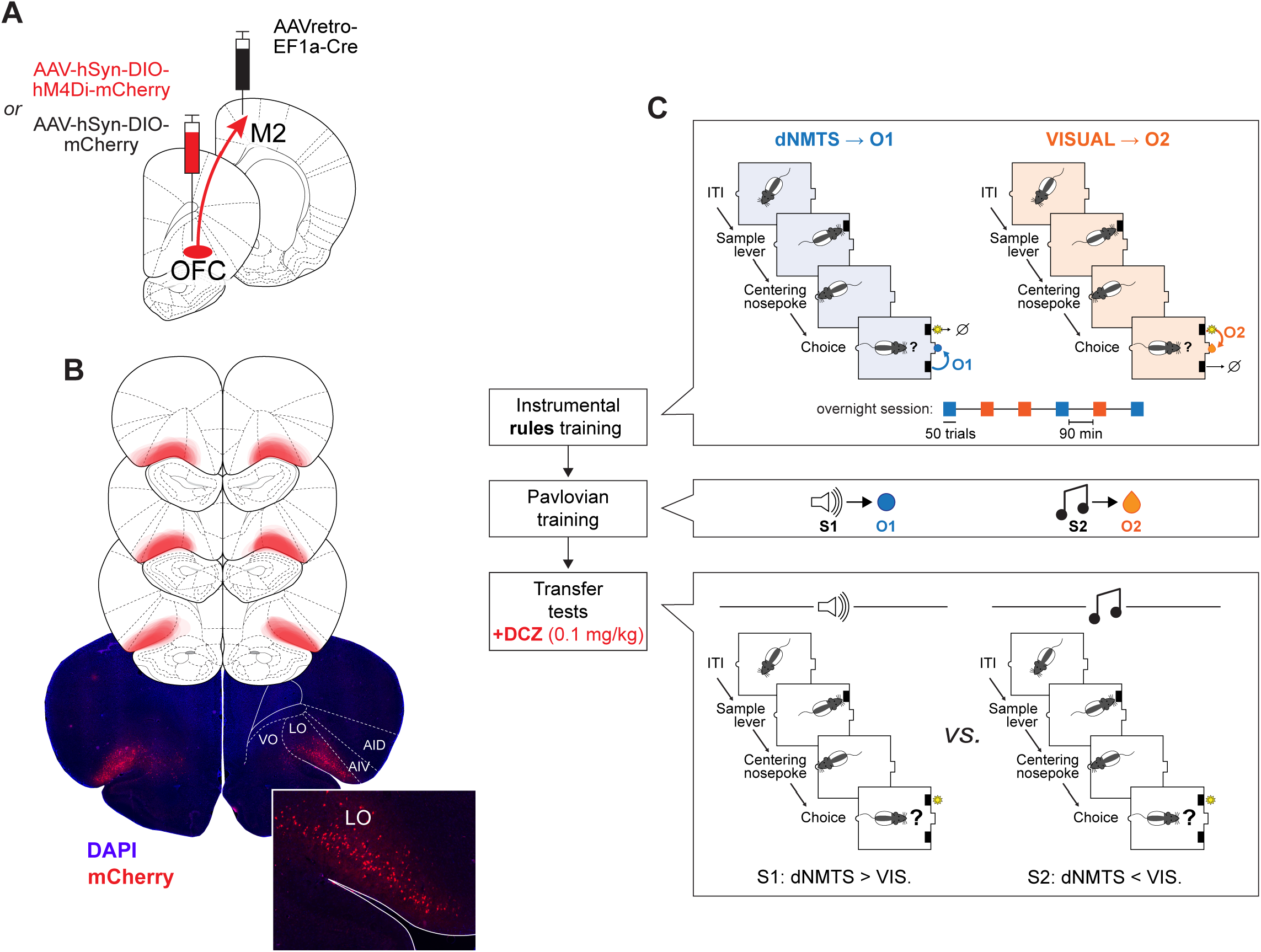
Pathway-specific chemogenetic inhibition of OFC→M2 neurons and Behavioral task design. **(A)** Intersectional viral strategy used for expressing the inhibitory DREADD (hM4Di-mCherry) or control transgene (mCherry) selectively in OFC neurons projecting to secondary motor cortex (M2). **(B)** Top: reconstructed representation of viral expression in the OFC (mCherry and hM4Di-mCherry subjects combined). Bottom: representative fluorescent image of mCherry expression in the OFC. **(C)**. Behavioral task design. Rats learned two instrumental rules governing completion of a three-step action sequence, with each rule producing a distinct outcome (dNMTS–O1; VISUAL–O2). In a second phase, two auditory stimuli were established as Pavlovian predictors of these outcomes (S1–O1; S2–O2). Finally, nonreinforced transfer tests assessed the influence of these Pavlovian cues on instrumental rule selection. All rats received DCZ (0.1 mg/kg, i.p.) 40 min before each transfer test. See methods for details.

In an initial phase, rats were gradually trained to complete a three-action instrumental sequence (sample lever → nosepoke → choice lever) to obtain food reward. Critically, across different training blocks, animals learned that two distinct abstract rules could be used to guide selection of the final choice lever. Successful performance under each rule was reinforced with a distinct reward outcome (banana pellet or chocolate milk; counterbalanced). Specifically, during delayed non-match-to-sample (dNMTS) blocks, rats learned to select the non-sampled lever to obtain outcome 1 (dNMTS–O1). Conversely, during visual-rule (VIS) blocks, rats learned to select the visually cued lever to obtain outcome 2 (VIS–O2) (**Figure 1C**). Training proceeded gradually according to the curriculum detailed in **Supplemental Table 1**.

By the end of instrumental rule acquisition and throughout the retraining period, rats completed overnight sessions in which dNMTS and VIS blocks alternated in a pseudorandom order (50 trials/block; inter-trial interval: 45 s; inter-block interval: 90 min). At the start of each block, a free reward (O1 or O2) was delivered to signal the identity of the upcoming block. Animals displayed high levels of accuracy on both rules (proportion correct > 90%). A two-way mixed-model ANOVA on choice accuracy revealed a significant main effect of Rule (F(1,20) = 6.832, P = 0.017), as rats performed slightly better on the VIS rule (mean accuracy = 94.6%) relative to the dNMTS rule (mean accuracy = 91.6%). Importantly, no main effect of Virus or Rule * Virus interaction was observed (all Ps ≥ 0.398), indicating similar levels of instrumental performance across groups (**Figure 2A**). Representative videos illustrating performance of the dNMTS and VIS rules are provided in Supplementary Video 1.

**Figure 2.**
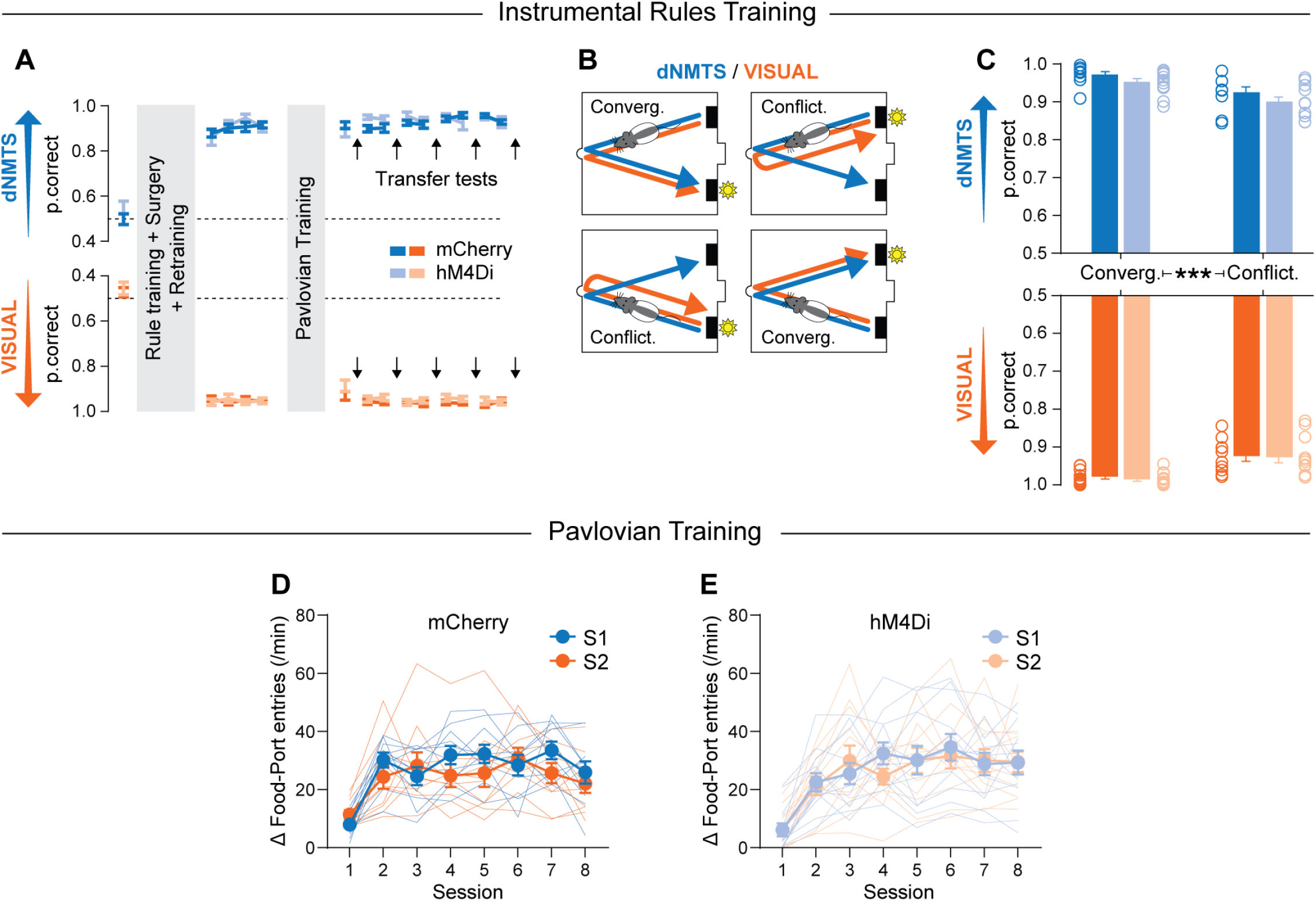
Acquisition of instrumental rules and Pavlovian cue-outcome associations. **(A)** Accuracy across instrumental rule training for the delayed non-match-to-sample (dNMTS; blue) and visual (orange) rules in mCherry and hM4Di groups. Arrows indicate transfer tests conducted after Pavlovian training. **(B)** Convergent trials required the same choice under both rules, whereas conflict trials required opposing choices. **(C)** Accuracy on convergent and conflict trials during the final three instrumental sessions preceding the first cue-guided transfer test. Performance was lower on conflict than convergent trials but did not differ between mCherry and hM4Di groups. **(D–E)** Cue-evoked food-port entry rates across Pavlovian training in the mCherry **(D)** and hM4Di **(E)** groups. Cue-evoked responding was calculated as the food-port entry rate during the cue, before the first reward delivery, minus the prestimulus entry rate. Data are presented as trial-averaged, between-subject mean ± s.e.m. Circles in **C** and thin lines in **D-E** represent individual data points. *** P < 0.001 post-hoc paired t test.

Because dNMTS and VIS rules were orthogonal, correct choice on certain trials could satisfy both rules simultaneously. We refer to these as *converging trials*, because both rules converged on the same choice lever. This contrasts with *conflicting trials* in which the two rules would lead to different choice responses (**Figure 2B**). Performance was analyzed using a three-way mixed-model ANOVA (Rule × Conflict × Virus). As expected, accuracy was slightly lower on conflicting trials (Conflict: F(1,20) = 105.556, P < 0.001), presumably reflecting increased response conflict on those trials. This effect similarly impacted performance across both rules and virus groups, as no significant interactions involving Conflict were observed (all Ps ≥ 0.191). Importantly, even on conflicting trials, performance remained highly accurate (VIS: 91.9%; dNMTS: 88.6%), confirming that all rats successfully acquired both instrumental rules (**Figure 2C**).

In a second phase, rats underwent Pavlovian conditioning in which two 2-min auditory stimuli (clicker and white noise, counterbalanced as S1 and S2) were each paired with intermittent delivery of one of the two reward outcomes (S1→O1, S2→O2). This procedure has been shown to promote the formation of sensory-specific stimulus–outcome associations (Lichtenberg et al., 2017; Sias et al., 2021). Across eight conditioning sessions, rats progressively increased anticipatory goal-approach responding during cue presentation (food-port entries measured after cue onset and before reward delivery; main effect of Session: F(4.111, 90.251) = 41.291, P < 0.001). No main effects of Cue identity (S1 vs. S2) or Virus (hM4Di vs. mCherry), nor any interactions involving these factors, were observed (all Ps ≥ 0.095), indicating that all rats successfully acquired both cue–outcome associations (**Figure 2D-E**). Following Pavlovian training, rats were briefly retrained on the instrumental task during overnight sessions conducted both before and between transfer tests.

Finally, all rats underwent three types of transfer tests conducted across five nonreinforced probe sessions: (1) a baseline test controlling for spontaneous bias in rule selection in the absence of task-relevant information; (2) cue-evoked bias tests, in which S1 and S2 were presented continuously during separate probe sessions; and (3) outcome-evoked bias tests, in which O1 and O2 were delivered noncontingently during separate sessions. All probe tests were preceded by DCZ injection (0.1 mg/kg, i.p., 40 min before testing), resulting in selective inhibition of the OFC→M2 pathway in hM4Di-expressing rats. Each probe session consisted of 20 trials, all of which were conflicting trials (for which the dNMTS and VIS rules were mutually exclusive). For each animal and session, we calculated a rule-bias index defined as the difference between dNMTS and VIS choices divided by the total number of completed trials. Positive values reflected a bias toward the dNMTS rule, whereas negative values reflected a bias toward the VIS rule (**Figure 3A**).

**Figure 3.**
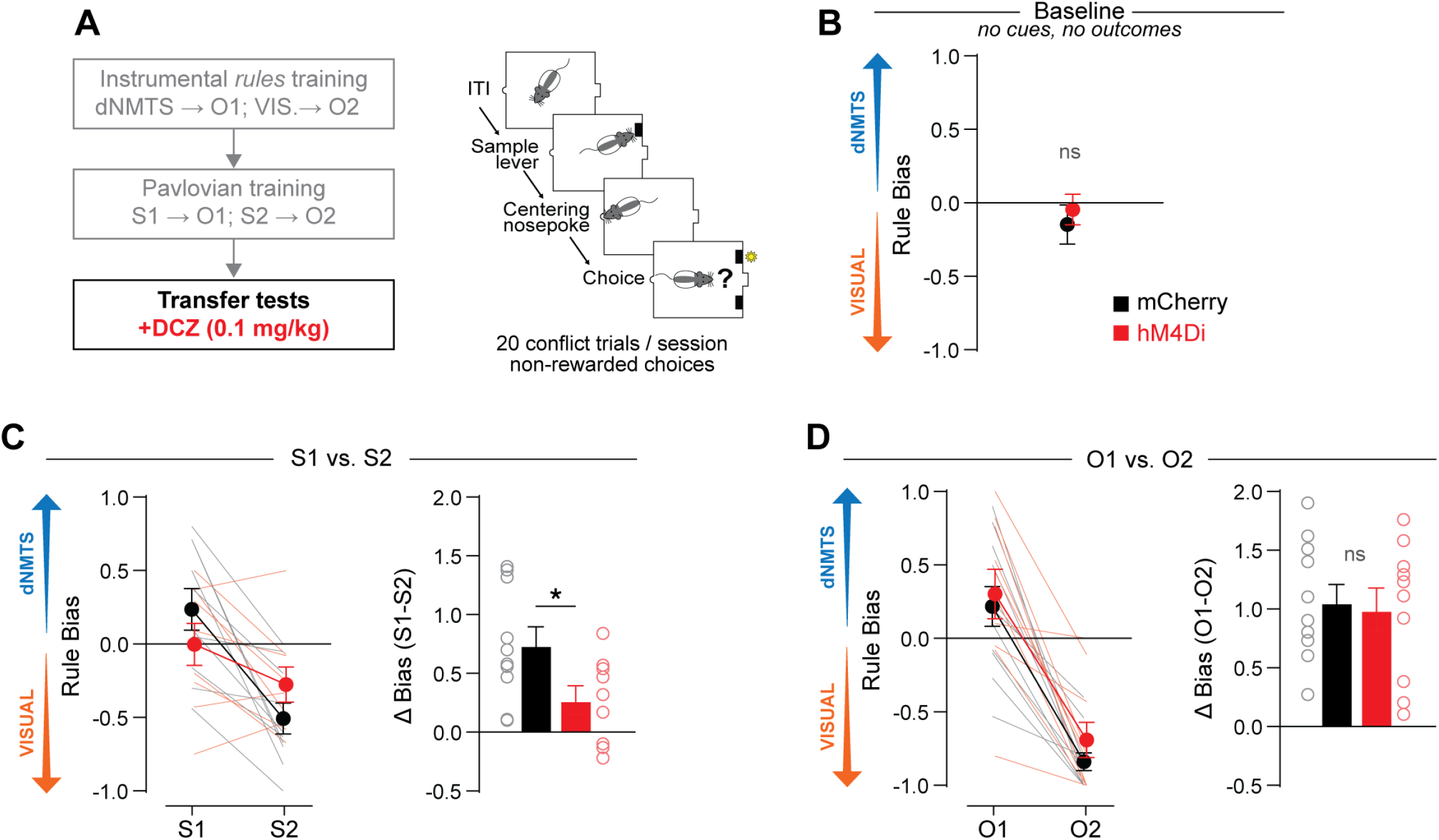
OFC→M2 inhibition disrupts cue-evoked, but not outcome-evoked, instrumental rule selection. **(A)** Overview of the transfer-test procedure. During each transfer test, rats completed 20 nonreinforced conflict trials. All rats received DCZ (0.1 mg/kg, i.p.) 40 min before each test. **(B)** Baseline test. In absence of cues or outcomes, neither group showed a spontaneous bias toward either instrumental rule. **(C)** Rule bias during cue-guided transfer tests. S1 biased rule selection toward dNMTS, whereas S2 biased selection toward the visual rule. This cue-induced bias (S1 − S2) was reduced by OFC→M2 inactivation in hM4Di rats relative to mCherry controls. **(D)** Rule bias during outcome-guided transfer tests. O1 biased rule selection toward dNMTS, whereas O2 biased selection toward the visual rule. This outcome-induced bias (O1 − O2) was spared by OFC→M2 inactivation. Data are presented as trial-averaged, between-subject mean ± s.e.m. Thin lines and open circles represent individual data points. * P < 0.05 post-hoc t test.

During the baseline test, rats displayed approximately equal numbers of dNMTS and VIS choices, indicating no spontaneous bias toward either strategy in the absence of task-relevant information (bias index not significantly different from zero; one-sample t test: t(21) = 0.508, p = 0.617). Importantly, OFC→M2 inhibition did not affect baseline rule selection bias (hM4Di vs. mCherry: t(20) = 0.023, p = 0.982) (**Figure 3B**). The total number of completed trials was also unaffected by OFC→M2 inhibition (mCherry: 18.18 ± 0.81; hM4Di: 17.09 ± 0.65; P = 0.306).

We next examined cue-evoked rule selection across two probe sessions (one cue per session). Pavlovian cues biased rule selection toward the corresponding strategy, such that S1 promoted dNMTS choices, whereas S2 promoted VIS choices (main effect of Cue: F(1,17) = 25.722, P < 0.001). Critically, this cue-driven bias was reduced by OFC→M2 inhibition, as evidenced by a Cue × Virus interaction (F(1,17) = 5.480, P = 0.032). Consistent with this, mCherry controls displayed a robust shift in rule bias between S1 and S2 sessions (P < 0.001), whereas hM4Di rats failed to show a significant difference in rule bias across cue conditions (P = 0.077) (**Figure 3C**). This impairment was not accompanied by a change in the total number of completed trials (mCherry: 17.15 ± 0.97; hM4Di: 15.22 ± 1.26; P = 0.236).

Finally, we examined the influence of the outcomes themselves on rule selection across two additional probe sessions (one outcome tested per session). The relevant outcome was delivered noncontingently 10 s before a trial, beginning with the first trial and every five trials thereafter. Because outcome delivery did not closely follow a choice response, this temporal arrangement minimized its direct reinforcing effects while promoting its discriminative, or cueing, influence on subsequent rule selection. As expected, outcome delivery biased rats toward the corresponding strategy, such that O1 promoted dNMTS choices whereas O2 promoted VIS choices (main effect of Outcome: F(1,19) = 76.382, P < 0.001). However, in contrast to cue-evoked transfer, this effect was unaffected by OFC→M2 inhibition, as indicated by the absence of main effects or interactions involving Virus (all Ps ≥ 0.397) (**Figure 3D**). Here again, OFC→M2 inhibition did not affect the total number of completed trials (mCherry: 17.45 ± 0.74; hM4Di: 15.75 ± 0.66; P = 0.103).

Together, these findings indicate that the OFC→M2 pathway is required for cue-evoked outcome memories —but not directly experienced outcomes— to bias abstract rule selection during reward pursuit.

### OFC→M2 inactivation does not impair performance on learned instrumental rules

Our results thus far indicate that OFC→M2 inactivation impairs reward-memory-guided rule selection. However, this effect could instead reflect an impairment in the ability to perform the instrumental rules themselves. We therefore examined whether OFC→M2 inactivation affected performance on the learned dNMTS and VIS rules under reinforced conditions. Following brief retraining, rats completed four test sessions in which performance on each rule (dNMTS or VIS) was assessed following treatment with either saline or DCZ (session order counterbalanced). Performance was analyzed using a mixed-model ANOVA with Rule (dNMTS vs. VIS), Virus (hM4Di vs. mCherry), and Treatment (saline vs. DCZ) as factors. OFC→M2 inactivation did not impair performance on either rule, as no main effects of Treatment or Virus, nor any interactions involving these factors were observed (accuracy > 86% in all groups and sessions; all Ps > 0.111). Restricting the analysis to conflicting trials yielded similar results, with no main effects or interaction of Treatment and Virus (accuracy > 79% in all groups and sessions; all Ps > 0.122).

These findings indicate that OFC→M2 inhibition does not impair execution of well-learned instrumental rules under reinforced conditions, supporting a more selective role for this pathway in using cued outcome memories to guide rule selection.

### OFC→M2 inactivation does not impair cue-evoked action selection

Our results thus far indicate that the OFC→M2 pathway is required for cue-evoked reward memories to bias abstract rule selection. We next sought to determine whether this pathway is specifically involved in abstract rule selection or instead contributes more broadly to reward-memory-guided behavior. To address this question, we tested the effects of OFC→M2 inhibition in a separate cohort of rats trained in a simpler task analogous to outcome-specific Pavlovian-to-instrumental transfer (sPIT). In this task, rats learned direct associations between discrete instrumental actions and distinct reward outcomes (right lever press→O1; left lever press→O2). We then examined whether Pavlovian cues evoking memories of O1 or O2 would bias action selection toward the corresponding lever, and whether this effect was disrupted by OFC→M2 inactivation.

As in the previous experiment, rats underwent three types of nonreinforced transfer tests: a baseline test, cue-evoked bias tests, and outcome-evoked bias tests. During the baseline test, rats displayed no spontaneous preference for either action in the absence of task-relevant information (lever bias index not significantly different from zero; one-sample t test: t(20) = 0.641, p = 0.529), and OFC→M2 inhibition had no effect on baseline action bias (p = 0.735) (**Figure 4A**).

**Figure 4.**
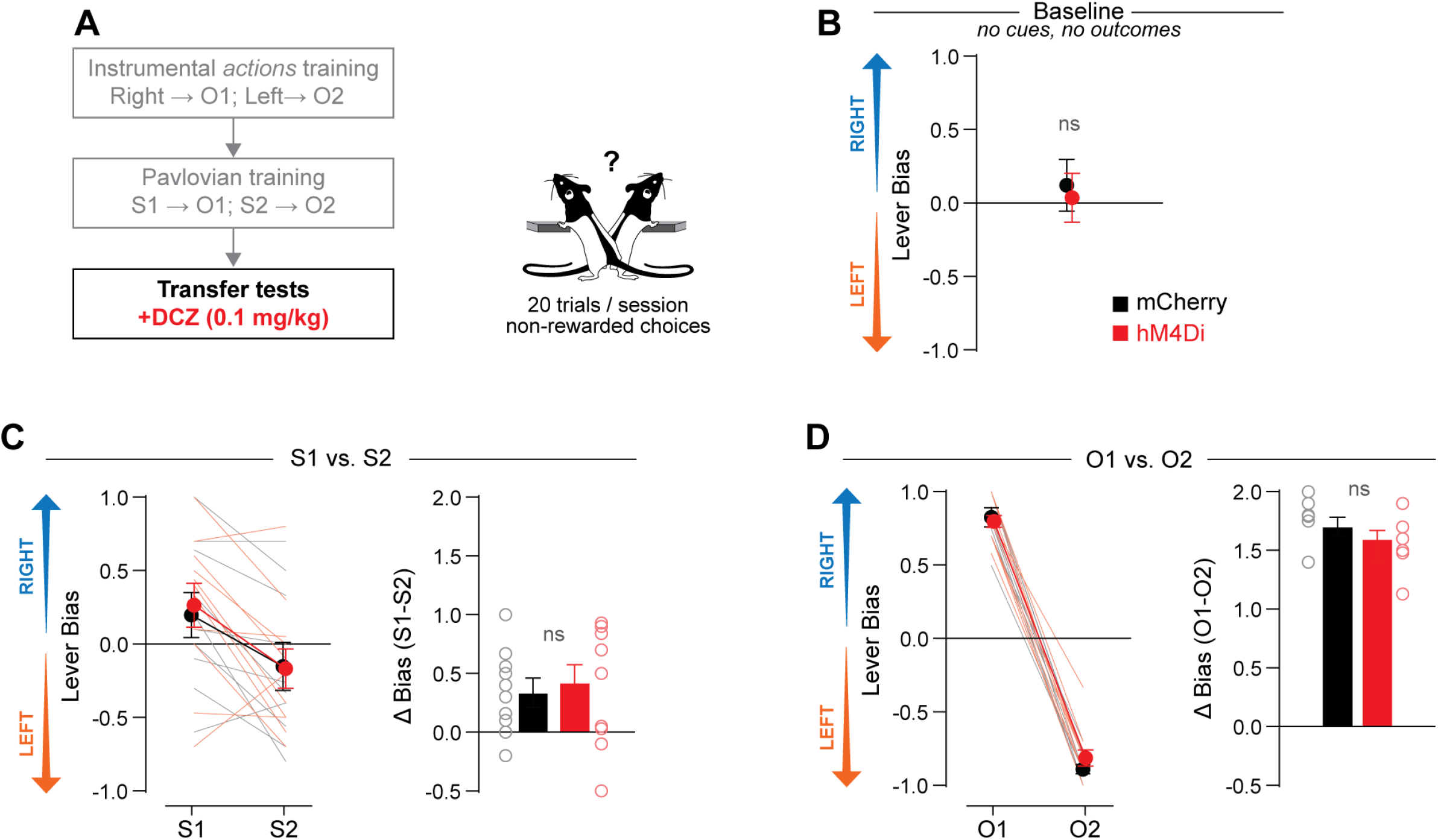
OFC→M2 inhibition spares cue- and outcome-evoked instrumental action selection. **(A)** Overview of the training and transfer-test procedure. Rats first received discrete action–outcome training (right–O1; left–O2), followed by Pavlovian training (S1–O1; S2–O2). During each transfer test, rats completed 20 nonreinforced choices between the two actions. All rats received DCZ (0.1 mg/kg, i.p.) 40 min before each test. **(B)** Baseline test. In the absence of cues or outcomes, neither group showed a spontaneous bias toward either instrumental action. **(C)** Lever bias during cue-guided transfer tests. S1 biased action selection toward the right lever, whereas S2 biased selection toward the left lever. This cue-induced bias (S1 − S2) was spared by OFC→M2 inactivation. **(D)** Lever bias during outcome-guided transfer tests. O1 biased action selection toward the right lever, whereas O2 biased selection toward the left lever. This outcome-induced bias (O1 − O2) was spared by OFC→M2 inactivation. Data are presented as trial-averaged, between-subject means ± s.e.m. Thin lines and open circles represent individual data points.

We next examined cue-evoked action selection across two probe sessions (one cue per session). As expected, Pavlovian cues biased responding toward the action associated with the corresponding reward outcome (main effect of Cue: F(1,19) = 17.713, p < 0.001). In contrast to the abstract rule task, however, this cue-evoked action bias was unaffected by OFC→M2 inhibition, with no main effect of Virus or Cue × Virus interaction (all Ps ≥ 0.660) (**Figure 4B**).

Finally, we examined the influence of noncontingent reward delivery on action selection. Outcome delivery strongly biased responding toward the corresponding instrumental action (main effect of Outcome: F(1,19) = 1289.382, P < 0.001), with rats selecting the corresponding lever on nearly every trial. This outcome-evoked action bias was similarly unaffected by OFC→M2 inhibition, with no main effect of Virus or Outcome × Virus interaction (all Ps ≥ 0.275) (**Figure 4C**).

Together, these findings indicate that the OFC→M2 pathway is selectively required for cue-evoked reward memories to influence abstract rule selection, but not the selection of discrete instrumental actions.

## DISCUSSION

Reward-predictive cues (i.e. Pavlovian cues) can influence instrumental behavior by activating representations of specific outcomes, thereby promoting the actions associated with those outcomes (Balleine and Ostlund, 2007; de Wit and Dickinson, 2009). This interaction between Pavlovian predictions and instrumental behavior is commonly studied using specific Pavlovian-to-Instrumental Transfer (sPIT) paradigms, in which a Pavlovian cue biases choice between two discrete actions (often a left or right lever press). However, in naturalistic reward seeking, individual actions are often organized into extended sequences, with higher-order rules determining which information is relevant and how it should guide action selection within those sequences. Here, we developed a new task —the specific Pavlovian-to-Rules-to-Instrumental Transfer (sPRInT) paradigm— to capture this intermediate level of organization between outcome representation and action selection. Rather than associating each outcome with a discrete action, rats learned that two different rules could govern completion of a common three-steps instrumental sequence, with each rule producing a distinct outcome. Because either rule could lead to a left or right choice depending on the information presented on a given trial, rule selection could be dissociated from selection of a particular motor response. This design allowed us to examine whether Pavlovian outcome predictions determine not only which action is selected, but also the higher-order rule that governs how ongoing information is transformed into action.

Using this paradigm, we found that Pavlovian cues biased rule selection, promoting selection of the rule corresponding to the predicted outcome. During nonreinforced probe sessions, S1 promoted use of the dNMTS rule (shared outcome: O1), whereas S2 promoted use of the VIS rule (shared outcome: O2). Chemogenetic silencing of OFC→M2 projecting neurons reduced this cue-evoked rule bias without affecting baseline rule preference or the number of completed trials. Moreover, OFC→M2 inactivation spared rule selection when outcome-specific information was provided by noncontingent delivery of the outcomes themselves, as well as accurate execution of both rules during reinforced sessions. Finally, in a separate cohort trained on a simpler task analogous to conventional outcome-specific PIT, OFC→M2 inactivation did not disrupt the ability of Pavlovian cues to bias selection between discrete instrumental actions.

Collectively, these result help delineate the contribution of OFC→M2 projecting neurons. The preserved cue-evoked action-selection effect in the sPIT paradigm indicates that rats could still retrieve outcome-specific Pavlovian memories and use them to guide selection between discrete instrumental actions. Moreover, preserved performance on both the dNMTS and VIS rules during reinforced tests indicates that rats retained knowledge of the rules and remained capable of using sampling history or visual information to select the appropriate response. The preserved bias produced by direct outcome delivery further indicates that the learned relationships between outcomes and rules remained behaviorally accessible. The deficit caused by OFC→M2 inactivation therefore emerged specifically when an outcome representation had to be retrieved from memory and used to select the corresponding rule. These findings suggest that OFC→M2-projecting neurons are critical for transforming cue-evoked outcome representations into the selection of higher-order instrumental rules.

This interpretation is consistent with the known functions of OFC and M2. The OFC is critical for encoding predictive relationships between events and generating sensory-specific representations of anticipated outcomes, allowing expected outcome identity to guide behavior (Rolls, 2004; McDannald et al., 2011; Stalnaker et al., 2014; Wilson et al., 2014; Howard et al., 2015; Costa et al., 2023). M2 on the other hand has been implicated in flexible action selection, movement planning, and the organization of complex action sequences (Sul et al., 2011; Gremel and Costa, 2013; Barthas and Kwan, 2017; Ohbayashi, 2021; Sánchez-Fuentes et al., 2021; Schreiner et al., 2022; Augusto et al., 2025). Consistent with a role in the higher-order organization of behavior, M2 lesions can spare simple instrumental responding while disrupting the organization and regulation of goal-directed sequences (Ostlund et al., 2009; Gremel and Costa, 2013). More generally, M2 has been proposed to organize sequential behavior by linking antecedent conditions —including external stimuli and preceding actions— to subsequent motor output (Barthas and Kwan, 2017). This transformation may be implemented through attractor dynamics, with distinct M2 activity states specifying how antecedent information is translated into motor output (Inagaki et al., 2019; Recanatesi et al., 2022). Distinct instrumental rules may therefore correspond to distinct attractor states that configure different mappings between task information and action. By signaling the identity of the anticipated outcome, OFC input could bias M2 activity toward the appropriate rule state, thereby determining which information (sampling history or visual information) guides the final choice in the sequence. In this framework, silencing OFC→M2-projecting neurons would prevent cue-evoked outcome representations from biasing M2 toward the appropriate rule state. Directly experienced outcomes, however, could continue to recruit that state through alternative routes conveying orosensory or consummatory motor signals (Barthas and Kwan, 2017).

The trial-based structure of the present paradigm introduces another important distinction from conventional PIT procedures. Whereas in free-operant PIT paradigms Pavlovian cues can both invigorate and bias instrumental responding (Corbit and Balleine, 2005), our sPRInT task primarily captures their biasing influence. Indeed, in our task, insertion of the sample and choice levers prompted initiation and completion of the instrumental sequence, and rats completed most trials even in the absence of Pavlovian cues, limiting the task’s sensitivity to cue-induced invigoration. Nevertheless, OFC→M2 inactivation did not affect the number of completed trials, indicating that its effect on rule selection was not accompanied by a detectable reduction in task engagement or sequence completion. However, because the task was not designed to measure cue-induced changes in response vigor, these findings do not exclude a contribution of OFC→M2 to Pavlovian invigoration under free-operant conditions.

One limitation of our projection-specific chemogenetic approach is that systemic DCZ inhibited OFC→M2-projecting neurons globally, suppressing their output to M2 but also to any potential collateral targets. However, single-neuron reconstructions indicate that superficial OFC neurons —where most of our labeled neurons were located— exhibit relatively simple projection patterns, with a substantial subset projecting exclusively to motor cortices (Wang et al., 2020). This anatomical organization may limit the contribution of alternative OFC output pathways to the observed effects. Nevertheless, future experiments using terminal-specific inhibition within M2 will help confirm the specific contribution of the OFC→M2 pathway to the effects reported here.

In conclusion, the sPRInT paradigm provides a new approach for examining how cue-evoked outcome memories influence the selection of higher-order (“abstract”) rules that organize action sequences during reward pursuit. Using this paradigm, we found that Pavlovian cues promoted adoption of the rule associated with the anticipated outcome and that this effect depended on the functional integrity of OFC→M2-projecting neurons. These findings expand the established role of reward-predictive cues from biasing discrete actions to configuring the higher-order rules that organize extended behavior, a process that may contribute to both adaptive and maladaptive reward seeking (Garbusow et al., 2022). More broadly, incorporating higher-order rules as a dimension of instrumental behavior may reveal contributions of neural circuits that remain undetected in conventional action-based paradigms.

## MATERIALS AND METHODS

### Subjects

Male and female Long–Evans rats (Charles River; 8 weeks old on arrival) were housed in same-sex pairs. Rats were maintained on a 12-h light/dark cycle (lights on from 8:00 AM to 8:00 PM). Overnight training sessions were conducted from 7:00 PM to 7:00 AM; all other behavioral sessions were conducted between 5:00 and 7:00 PM. Except during pre- and postoperative recovery periods, when food was available ad libitum, rats were mildly food restricted to maintain approximately 95% of age-matched free-feeding weights. All experimental procedures were approved by the University of California, Santa Barbara Institutional Animal Care and Use Committee and conducted in accordance with National Institutes of Health guidelines.

### Stereotaxic Surgeries

Rats were anesthetized with isoflurane (4% induction; 1–2.5% maintenance) and prepared for aseptic surgery. Under stereotaxic guidance, a retrograde viral vector encoding Cre recombinase (Addgene #55636-AAVrg) was injected bilaterally into the secondary motor cortex (M2), resulting in Cre expression in M2-projecting neurons, including those originating in the orbitofrontal cortex (OFC). During the same surgery, rats received bilateral OFC injections of either AAV8-hSyn-DIO-hM4Di-mCherry (Addgene #44362-AAV8) or the control vector AAV8-hSyn-DIO-mCherry (Addgene #50459-AAV8). This intersectional viral strategy restricted expression of the inhibitory DREADD hM4Di, or the control transgene mCherry, to OFC neurons projecting to M2.

For M2 injections, 0.8 μL of virus was delivered at each of the following coordinates relative to bregma and the skull surface (in mm): AP, +2.3 and +0.5; ML, ±1.0; and DV, −2.9. For OFC injections, 0.8 μL of virus was delivered at each of the following coordinates: AP, +3.8; ML, ±2.6; and DV, −5.8 and −5.4. Viruses were delivered through 33-gauge needles (Hamilton, #65460-03) connected to an infusion pump at a rate of 1 μL/min. After each infusion, the needle remained in place for an additional 5 min to permit viral diffusion. Rats recovered for at least 1 week before behavioral training resumed. Chemogenetic silencing with deschloroclozapine (DCZ) was conducted at least 5 weeks after surgery.

### Apparatus and Rewards

Behavioral training was conducted in 12 identical conditioning chambers, enclosed in individual sound-attenuating cubicles (Med Associates). A fan mounted on the cubicle provided ventilation and low background noise. Two ceiling-facing lights located on the front and back internal walls of the cubicle provided diffuse chamber illumination. The front panel contained a centrally positioned recessed food port flanked by two retractable levers, with a cue light located above each lever. The back panel contained an internally illuminated nose-poke operandum. Two auditory stimuli —a clicker and a white-noise generator— were located on the front and rear panels, respectively (approximately 76 dB each). Each chamber was also equipped with a syringe pump, located outside the sound-attenuating cubicle, and a pellet dispenser for delivering liquid and solid rewards, respectively, into the food port. Experimental events and data collection were controlled by a computer running Med-PC software (Med Associates).

The reward outcomes consisted of 0.17 mL of chocolate milk (2:1 Nesquik® in water + 5% sugar, w/v; delivered over 3 s) and a 45-mg banana-flavored pellet (Bio-Serv, Flemington, NJ; formula F0059). Assignment of these rewards as O1 and O2 was counterbalanced across rats. The two outcomes were matched for caloric content and, based on pilot experiments, were equally preferred on average.

### Specific Pavlovian-to-Rules-to-Instrumental Transfer (sPRInT)

#### Phase I: Instrumental rules training

Rats were gradually trained to complete a three-step instrumental sequence (sample lever press → nose poke → choice lever press) to earn a reward. The training curriculum is detailed in **Supplemental Table 1**. During the final stages of training, each trial began with the extension of a randomly selected sample lever (left or right). A press on this lever within 30s caused it to retract and illuminated the nose-poke operandum on the rear panel. A nose poke within 20s extinguished the nose-poke light and extended both levers on the front panel. Simultaneously, the cue light above one randomly selected lever was illuminated. Rats then had 20 s to make a choice by pressing one of the two levers; correct choices resulted in reward delivery. Both levers were retracted at the end of each trial. Failure to complete any step within the allotted time terminated the trial and was recorded as an omission.

Critically, rats learned two abstract rules for completing the sequence, each associated with a distinct outcome. During delayed non-match-to-sample (dNMTS) blocks, rats were required to select the non-sampled lever (and disregard the lever cue light) to obtain outcome 1 (dNMTS–O1). During visual (VIS) blocks, rats were required to select the lever indicated by the cue light (and disregard previous sampling information), to earn O2 (VIS–O2).

Rats were initially trained on the two rules in separate sessions. They subsequently completed overnight sessions comprising six blocks —three dNMTS and three VIS— presented in a randomized order. Each block consisted of 50 trials separated by a 45-s intertrial interval (ITI). The beginning of each block was signaled by illumination of the house light and one noncontingent delivery of the outcome associated with that block, indicating both the available outcome and the rule in effect. Consecutive blocks were separated by 90-min intervals during which the house light remained off. Initially, all correct choices were rewarded. During the final stage of training, 90% of correct choices were rewarded to promote persistent responding under the nonreinforced conditions used during the subsequent transfer tests. Representative videos illustrating performance of the dNMTS and VIS rules are provided in Supplementary Video 1.

#### Phase II: Pavlovian training

Two auditory stimuli (white noise and clicker, counterbalanced as S1 and S2) were established as Pavlovian predictors of the two outcomes (S1–O1 and S2–O2) across eight sessions. Each session comprised 16 cue presentations (eight per cue) in a pseudorandom order. Each cue was presented for 2 min, with a mean ITI of 8 min. During cue presentation, the corresponding outcome was delivered on a 30s random time schedule, resulting in an average of four outcome deliveries per cue presentation.

#### Instrumental Rule reminder

Following Pavlovian training, rats received one overnight reminder session on the two instrumental rules (as described above) before the first transfer test. Additional reminder sessions were conducted between subsequent transfer tests (two overnight reminder between each transfer test).

#### Phase III: Transfer tests

During transfer tests, rats were prompted to complete the three-step instrumental sequence (sample-lever press → nose poke → choice-lever press). Each test comprised 20 nonreinforced rule-choice trials, organized into four series of five trials separated by 5-min intervals. Only conflict trials (for which dNMTS and VIS rules specified different choice levers) were presented, allowing each choice to be classified as consistent with one rule or the other. All rats received deschloroclozapine (DCZ; 0.1 mg/kg, i.p.) 40 min before testing. Three types of transfer tests were conducted across five sessions:

##### *Baseline* (session 1)

Neither Pavlovian cues nor outcomes were presented. This test assessed spontaneous bias in rule selection.

##### *Cue-induced rule-selection* (sessions 2 and 3)

Either S1 or S2 was presented beginning 10 s before the first trial of each five-trial series and remained on throughout the series. S1 and S2 were tested in separate sessions, with test order counterbalanced across rats.

##### *Outcome-induced rule-selection* (sessions 4 and 5)

Either O1 or O2 was delivered noncontingently 10 s before the first trial of each five-trial series. O1 and O2 were tested in separate sessions, with test order counterbalanced across rats.

### Specific Pavlovian-to-Rules-to-Instrumental Transfer (sPRInT)

#### Phase I: Instrumental action training

Unlike in the sPRInT task, in which outcomes were associated with abstract rules governing a three-step action sequence, in this task rats were trained to associate each outcome with a discrete instrumental action. Right- and left-lever presses earned two distinct outcomes (right–O1 and left–O2), with pellet and chocolate milk counterbalanced as O1 and O2 across rats.

Each trial began with the extension of both levers, and the block in effect determined which action was rewarded. During right-action blocks, right-lever presses earned O1 (right–O1), whereas left-lever presses were not rewarded. During left-action blocks, left-lever presses earned O2 (left–O2), whereas right-lever presses were not rewarded. Both levers retracted after a choice. Failure to choose a lever within 20 s terminated the trial and was recorded as an omission. As in sPRInT, rats ultimately completed overnight sessions comprising six 50-trial blocks (three blocks of each type) presented in a pseudorandom order. Within a block, trials were separated by a 45-s ITI, and consecutive blocks were separated by 90-min intervals during which the house light remained off. Each block began with house-light illumination and one noncontingent delivery of the associated outcome, indicating the action rewarded during that block. Initially, all correct choices were rewarded. During the final stage of training, 90% of correct choices were rewarded to promote persistent responding under the nonreinforced conditions used during the subsequent transfer tests.

#### Phase II: Pavlovian training

Pavlovian training was conducted as described for the sPRInT procedure.

As in sPRInT, rats received one overnight reminder session before the first transfer test and two additional overnight reminder sessions before each subsequent test.

#### Phase III: Transfer tests

The three types of transfer tests—baseline, cue-induced, and outcome-induced—were conducted across five sessions as described for sPRInT, except that each trial consisted solely of a choice between the left and right levers.

### Histological verification of DREADD expression

Rats were deeply anesthetized with pentobarbital and transcardially perfused with phosphate-buffered saline (PBS), followed by 4% paraformaldehyde (PFA). Brains were extracted, postfixed in 4% PFA for 24 h, cryoprotected in 30% sucrose in PBS for at least 3 days, and coronally sectioned at 40 μm using a cryostat. Sections were mounted onto glass slides and coverslipped with a DAPI-containing mounting medium (EverBrite^TM^ #23004; Biotium). Viral mCherry expression in the OFC was examined using a fluorescence microscope (BZ-X800; Keyence).

### Statistics and analyses

For each rat trained in the sPRInT task, a rule-bias index was calculated as:

Rule bias index = (dNMTS choices-VIS choices) / (total choices)

Positive values indicate a bias toward the dNMTS rule, whereas negative values indicate a bias toward the VIS rule.

For rats trained in the sPIT task, an analogous lever-bias index was calculated as:

Lever bias index = (RIGHT lever choices-LEFT lever choices) / (total choices)

Positive and negative values indicate biases toward the left and right levers, respectively.

Statistical analyses were performed using IBM SPSS Statistics (version 28). Specific analyses are described throughout the Results section and compiled in **Supplemental Table 2**. Analyses generally consisted of mixed-design ANOVAs with Cue or Outcome as a within-subject factor and Virus (hM4Di or mCherry) as a between-subject factor. Greenhouse–Geisser corrections were applied when the assumption of sphericity was violated, resulting in noninteger degrees of freedom where applicable. Post hoc comparisons were conducted using Šidák-adjusted t tests. Significance was assessed against a type I error rate of 0.05. Effect sizes are reported as partial eta squared (ηp^2^) for ANOVA main effects and interactions, and Cohen’s d for post hoc comparisons.

Initial group sizes were sPRInT_mCherry n = 11; sPRInT_hM4Di n = 11; sPIT_mCherry n = 10; and sPIT_hM4Di n = 11. Rats that completed less than 8 trials in a transfer test were excluded from the corresponding statistical analysis. As a result, sample sizes for the baseline, cue-induced, and outcome-induced analyses were, respectively: sPRInT_mCherry n = 11, 10, and 11; sPRInT_hM4Di n = 11, 9, and 10; sPIT_mCherry n = 10, 10, and 10; and sPIT_hM4Di n = 11, 11, and 11.

No significant main effects or interactions involving Sex were detected; therefore, data from males and females were collapsed for analysis. However, the study was not powered to assess sex differences.

---

**Supplemental Table 1.** Instrumental rules-training curriculum. *Remedial trials: an incorrect choice resulted in the same trial being repeated. Following two consecutive errors, the trial was repeated as a forced-choice trial, in which only the correct final lever was extended. Forced-choice trials were excluded from accuracy calculations.

| Stage | Procedure | Description | Outcome and reinforcement | Criterion / Training duration |
| --- | --- | --- | --- | --- |
| 1 | magazine training | 60 sucrose deliveries at ~1-min intervals | Sucrose; noncontingent | 2 sessions |
| 2 | lever training | Both levers available concurrently; each retracted after 30 reinforced presses | Sucrose; FR1 | 3–4 sessions; max. 2 h/ session (or 60 reinforcers) |
| 3 | nosepoke→lever training | Nosepoke initiated each trial and caused a lever to extend. Lever presses were reinforced on an FR1 schedule | Sucrose; FR1 | 3–4 sessions; max. 2 h/ session (or 60 reinforcers) |
| 4 | dNMTS forced choice no lever cue | 3-step sequence. At the final choice step, only the correct (nonsampled) lever was presented. No lever cue light was illuminated. | O1; FR1 | 3 sessions; 50 trials/session |
| 5 | dNMTS free choice no lever cue | 3-step sequence. At the final choice step, both levers were presented. No lever cue light was illuminated. Selection of the nonsampled lever was reinforced. | O1<br>P(rew.\|correct) = 1<br>Remedial trials*: Y | 8-12 sessions; Criterion: ≥ 85% accuracy |
| 6 | VIS free choice | At the final choice step, both levers were presented and one lever cue was illuminated. Selection of the visually cued lever was reinforced. | O2<br>P(rew.\|correct) = 1<br>Remedial trials*: Y | 8-12 sessions; Criterion: ≥ 85% accuracy |
| **Surgery and recovery** |  |  |  |  |
| 7 | dNMTS free choice no lever cue | Post-operative reminder of stage 5 | O1<br>P(rew.\|correct) = 1<br>Remedial trials*: Y | 2-4 sessions; Criterion: ≥ 85% accuracy |
| 8 | VIS free choice | Post-operative reminder of stage 6 | O2<br>P(rew.\|correct) = 1<br>Remedial trials*: Y | 2-3 sessions; Criterion: ≥ 85% accuracy |
| 9 | Combined dNMTS (no cue) & VIS | Overnight sessions comprising six blocks (three dNMTS and three VIS blocks) presented in pseudorandom order. No lever cue during dNMTS blocks | O1 and O2<br>P(rew.\|correct) = 1<br>Remedial trials*: Y | 2 overnight sessions; Criterion: ≥ 85% accuracy |
| 10 | Combined dNMTS & VIS | Overnight sessions comprising six blocks (three dNMTS and three VIS blocks) presented in pseudorandom order. Lever cue presented on all trials (relevant during VIS and irrelevant during dNMTS blocks) | O1 and O2<br>P(rew.\|correct) = 1<br>Remedial trials*: Y | 4 overnight sessions; Criterion: ≥ 85% accuracy |
| 11 | Combined dNMTS & VIS | Same procedure as Stage 10, with partial reinforcement and no remedial trials | O1 and O2<br>P(rew.\|correct) = 0.9<br>Remedial trials*: N | 4-5 overnight sessions; Criterion: ≥ 85% accuracy |
| **Pavlovian training** |  |  |  |  |
| 12 | Combined dNMTS & VIS | Same procedure as Stage 11; conducted after Pavlovian training and between transfer testing | O1 and O2<br>P(rew.\|correct) = 0.9<br>Remedial trials*: N | 9 overnight sessions |

**Supplemental Table 2.**
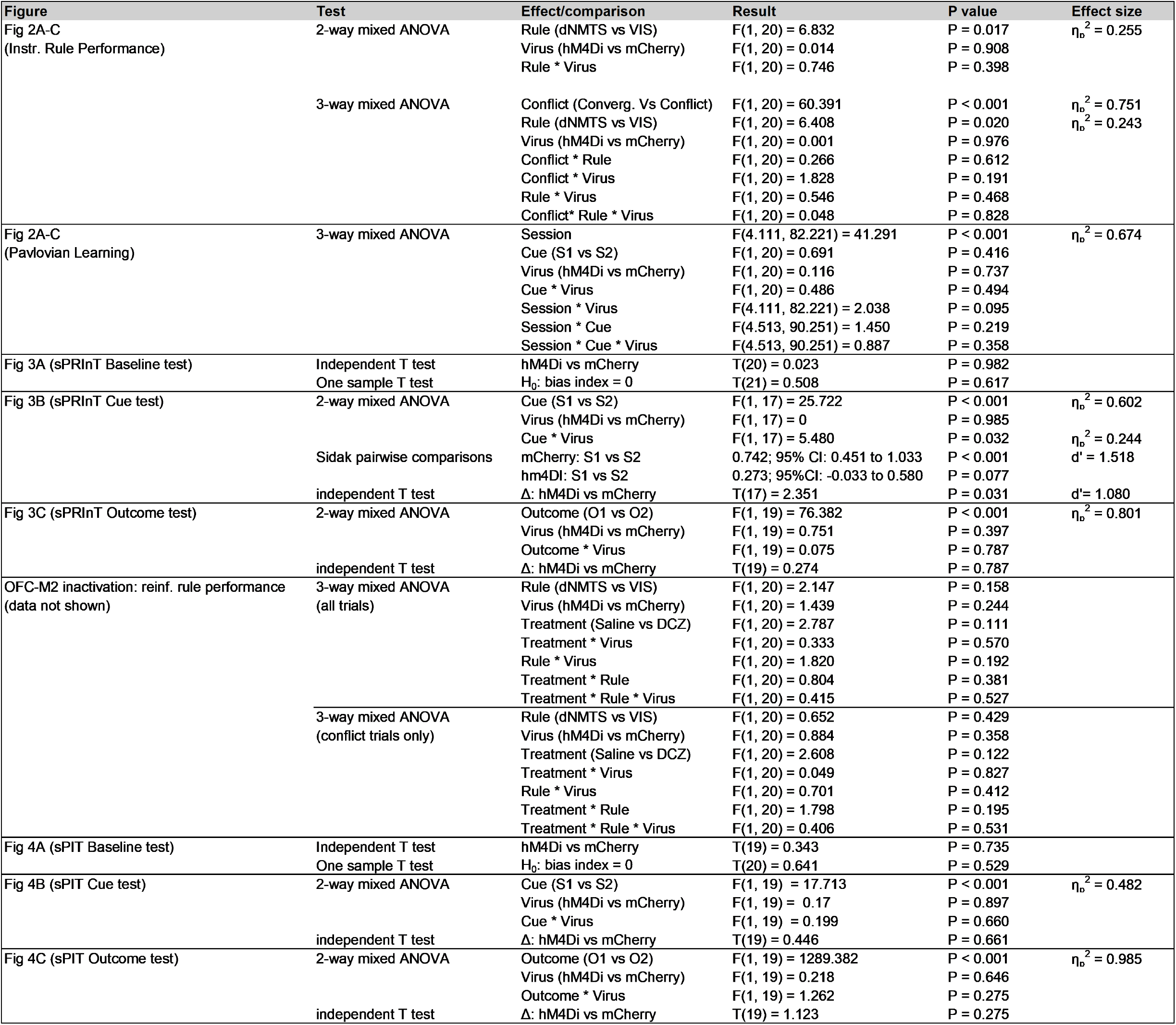
Comprehensive statistical results.

## Notes

### Competing Interest Statement

The authors have declared no competing interest.

